# Functional non-coding SNPs in human endothelial cells fine-map vascular trait associations

**DOI:** 10.1101/2021.08.03.454513

**Authors:** Anu Toropainen, Lindsey K. Stolze, Tiit Örd, Michael Whalen, Paula Martí Torrell, Verena M. Link, Minna U Kaikkonen, Casey Romanoski

## Abstract

Functional consequences of genetic variation in the non-coding human genome are difficult to ascertain despite demonstrated associations to common, complex disease traits. To elucidate properties of functional non-coding SNPs with effects in human endothelial cells (EC), we utilized molecular Quantitative Trait Locus (molQTL) analysis for transcription factor binding, chromatin accessibility, and H3K27 acetylation to nominate a set of likely functional non-coding SNPs. Together with information from genome-wide association studies for vascular disease traits, we tested the ability of 34,344 variants to perturb enhancer function in ECs using the highly multiplexed STARR-seq assay. Of these, 5,592 variants validated, whose enriched attributes included: 1) mutations to TF binding motifs for ETS or AP1 that are regulators of EC state, 2) location in accessible and H3K27ac-marked EC chromatin, and 3) molQTLs associations whereby alleles associate with differences in chromatin accessibility and TF binding across genetically diverse ECs. Next, using pro-inflammatory IL1B as an activator of cell state, we observed robust evidence (>50%) of context-specific SNP effects, underscoring the prevalence of non-coding gene-by-environment (GxE) effects. Lastly, using these cumulative data, we fine-mapped vascular disease loci and highlight evidence suggesting mechanisms by which non-coding SNPs at two loci affect risk for Pulse Pressure/Large Artery Stroke, and Abdominal Aortic Aneurysm through respective effects on transcriptional regulation of *POU4F1* and *LDAH*. Together, we highlight the attributes and context dependence of functional non-coding SNPs, and provide new mechanisms underlying vascular disease risk.

## Introduction

Genome-Wide Association Studies (GWAS) have revealed thousands of associations between genetic variants and clinical phenotypes. A large majority of disease associated loci, and thus functional variants that underpin trait differences, are not protein coding^1^. This suggests that non-coding disease-associated variants alter transcription through mechanisms such as recruitment of transcriptional activators and/or repressors at regulatory elements, or by changing chromatin conformation. Because regulatory elements, especially enhancers, are frequently cell-type specific^2^ or restricted by cell state in their activities^3; 4^, identification of disease-predisposing cells and tissues within the body remains a barrier toward functional understanding of risk loci.

Despite significant advancements in the catalogs linking human sequence variants to clinical phenotypes and molecular ‘omic profiles, a major bottleneck in functional genomics remains; namely, pinpointing *functional* regulatory variants and their mechanisms of action in relevant biological systems at scale. The identification of functional variants relative to proxy variants in high linkage disequilibrium (LD) presents an additional challenge. Recent studies have begun to map Quantitative Trait Loci (QTLs) for gene regulatory traits, such as histone modification, transcription factor (TF) binding, and chromatin accessibility^5^. These studies have identified allelic differences that associate with quantitative epigenetic differences in *cis*. However, causative roles for regulatory variants require experimental validation by separation from linked variants. To this end, Massively Parallel Reporter Assays (MPRAs) represent a high-throughput solution for experimental validation, compared to luciferase or EMSA assays, since several thousand sequences can be tested for regulatory function simultaneously^6-8^. Various MPRA techniques have been developed and implemented to identify genomic sequences and compare allele-specific effects on enhancer activity^8-13^.

In a recent study, we mapped QTLs for molecular traits in a set of genetically diverse Human Aortic Endothelial Cells (HAECs)^14^ under basal and pro-inflammatory conditions mimicked by cytokine Interleukin 1 beta (IL1B) stimulation. This system was chosen as a model for understanding mechanisms by which common genetic variants in humans shape molecular traits in vascular biology and inflammatory pathologies including atherosclerosis. Through that study, we identified thousands of *cis-*variants associated with transcript levels, chromatin accessibility, histone H3 lysine 27 acetylation (H3K27ac), and DNA binding of the transcription factors ERG and p65 (a component of NF-kB)^14^. This thoroughly phenotyped genetic panel of human cells provides a unique opportunity to pinpoint functional regulatory variants as well as to experimentally define genomic features that best correspond to validation in MPRAs. To this end, we extend the analysis of this resource by generating a matching MPRA dataset using Self-Transcribing Active Regulatory Regions with Sequencing (STARR-seq). This allowed us to investigate the correlation between allele-specific activity in STARR-Seq and molecular QTLs, while identifying attributes that best associate with variant effects. We demonstrate that allele-specific activity often depends on environmental stimulation (IL1B). Finally, we present several examples where our integrative approach allowed for functional fine-mapping of non-coding variants at cardiovascular GWAS loci.

## Results

### STARR-seq validates ETS and AP-1 factor motifs enriched in HAEC enhancer element

By overlaying evidence from GWAS, our molecular QTL (molQTL) and expression QTL (eQTL) study^14^, and allele-specific motif mutation analysis (see **Methods**), we selected 34,344 variants, represented by 59,977 oligos, for inclusion in a STARR-seq library (**Fig. S1)**. In STARR-seq, enhancer function was tested by placement of a ∼200 base pair oligo downstream of core promoter (origin of replication, ORI), followed by a poly-adenylation (poly-A) track, such that oligos that enhance transcription can be identified by sequencing of the produced non-coding RNAs (**Fig. 1A**). For all variants, both alleles were represented by oligos as they occur in haplotypes in the 1000 Genomes European reference population^15^ and surrounded by their native genomic sequences. The of STARR-seq libraries were transfected into teloHAECs (HAECs immortalized with hTERT) and enhancer activity was measured as the ratio of transcripts from each oligo relative to the amount of DNA plasmid for that oligo (i.e., RNA/DNA ratio). As expected, oligos that overlapped EC-specific enhancer elements had significantly greater STARR-seq activity compared to negative or scrambled oligo sequences (**Fig. 1B**). *De novo* motif analysis of the upper 10^th^ percentile of oligos with greatest enhancer activity were enriched for the JUN(AP-1), ETV2(ETS), and NFkB TF motifs when compared to all sequences in the STARR-seq library (**Fig. 1C**). This is consistent with our previous reports of these motifs being enriched at HAECs enhancers in the native chromatin context^5; 16^. More specifically, the ETS motif was frequently bound by the ERG in HAECs, which is an essential transcription factor (TF) regulating vascular development in mice^17^. Similarly, the AP-1 motif was also found as enriched at HAEC enhancers and bound by JUN^5^. These data provide evidence that enhancer activity measured by STARR-seq in teloHAECs, and by ChIP-seq in the genomic context of HAECs, are concordant. Therefore, the STARR-seq system is expected to reliability quantify the allelic effects of regulatory variants

**Figure 1.**
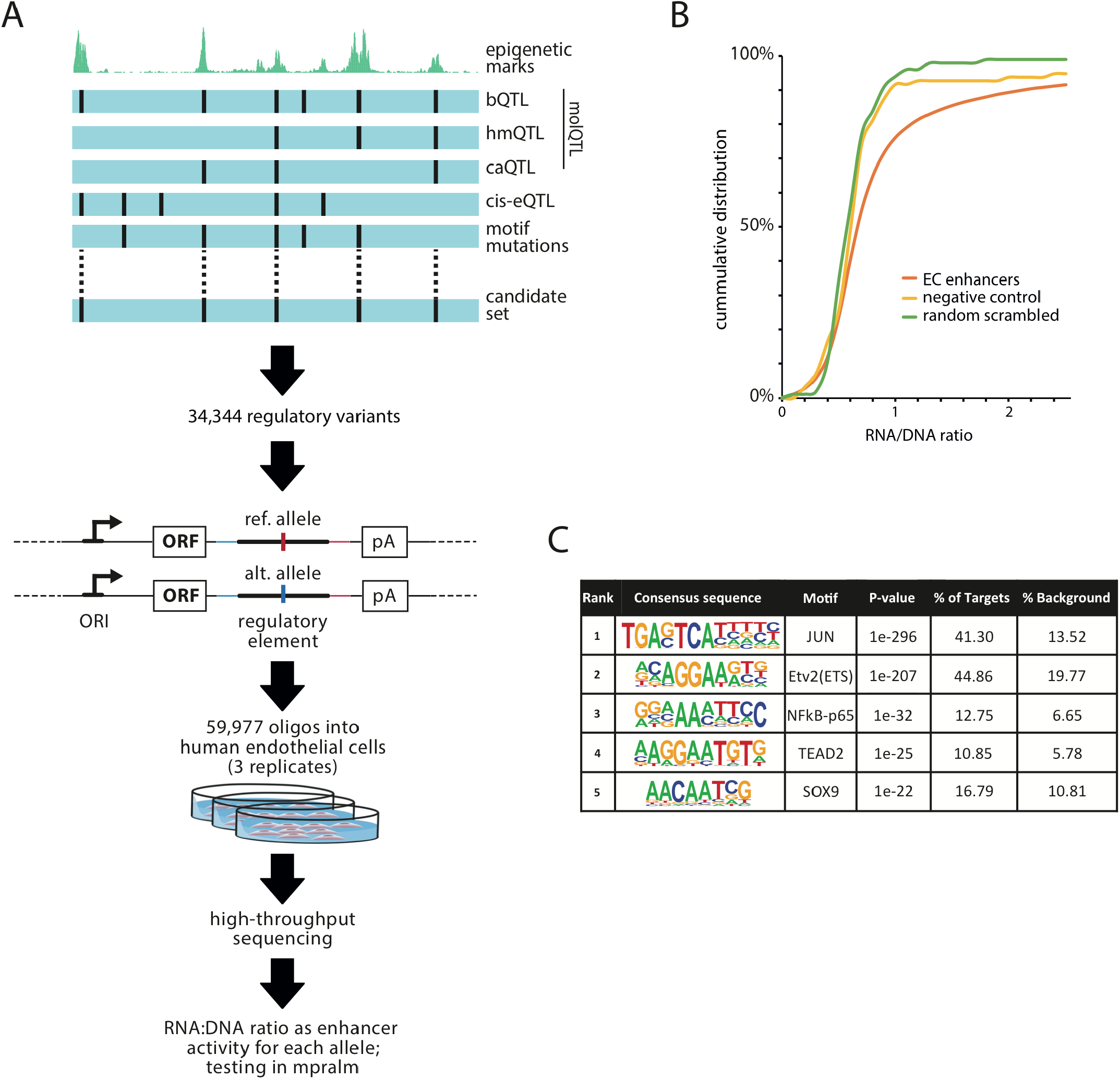
Construction and characterization of the STARR-Seq library. **A)** Schematic of the STARR-Seq library design. A candidate set of 198 bp oligos were selected based on their overlap with binding (b)QTLs, histone mark (hm)QTLs, chromaton accessibility (ca)QTLs and cis-eQTLs. Reference and alternative variants were cloned in plasmid vector under origin of replication core-promoter. The reporter library was transfected into cultured teloHAECs and deep sequencing was conducted. The enrichment of reporter RNA expression over input DNA library directly and quantitatively reflects enhancer activity. **B)** Cumulative histogram of enhancer activity for oligos overlapping active chromatin marks (H3K27ac, H3K4me2 and ATAC-Seq in ECs) compared to random regions and oligos outside active epigenetic elements. **C)** De novo motif analysis of the oligos showing enhancer activity within the upper 10th percentile using Homer.

### Among sequence attributes, TF motif mutations are most enriched for allele-specific regulatory activity

We next applied mpralm^18^ to analyze the allele-specific enhancer activity in the STARR-seq library and found that 3,256 variants (9.7% of those passing QC) were significant at 5% FDR and that triplicate biological replicates were highly concordant (**Fig. 2A**). For simplicity, these variants are referred to as ‘validated’ variants. To test whether sequence-based genomic features differentiated validated from un-validated variants, we evaluated the following metrics: 1) GC content in the STARR-seq oligos, 2) genomic distance of variants to the nearest transcription start sites (TSS), 3) average sequence conservation across vertebrates in oligo sequences, and, 4) SNPs that mutate TF binding motifs. Enrichment Scores (ES) were calculated in each analysis by performing a binomial test for significant differences between the number of validated SNPs in the category (eg. motif mutating SNPs) and the number of validated SNPs expected in that category based on how many were input into the library. Of the genomic features tested, we observed that oligos with GC content between 0.365 and 0.469 tended to harbor SNPs with allele-specific enhancer activity; similarly, regions with greater conservation scores were more likely to validate (**Fig. S2A**). Notably, regions with 60-70% GC started to be underrepresented in the library with a concurrent increase in data variability whereas GC>80% were rarely detected, suggesting that high GC content limits the detection accuracy (**Fig S2B**,**C)**. Still, only 2% of the input library had GC>70%, which is why we expect this to minimally affect downstream analysis. Additionally, TF motif mutating SNPs were significantly associated, as a set, with allele-specific regulatory activity (**Fig. 2G**); however, we reasoned this relationship was driven by a particular set of motifs corresponding to EC-lineage determining TFs. Among the TF motifs with strongest associations were members of the AP-1 family (Atf3, AP1, and Fra1), as well as members of the ETS family (ERG, ETS1, FLI1, ETV2) (**Fig. 2C-E**). Both AP-1/ATF3 and ETS/ERG motifs were enriched over random expectations qualitatively (ES scores), and quantitatively such that greater allelic effects on motif scores relate to more significant STARR-seq p-values (**Fig. 2D**,**E**). These data are consistent with AP-1 and ETS interactions at these loci being important for HAEC regulatory function.

**Figure 2.**
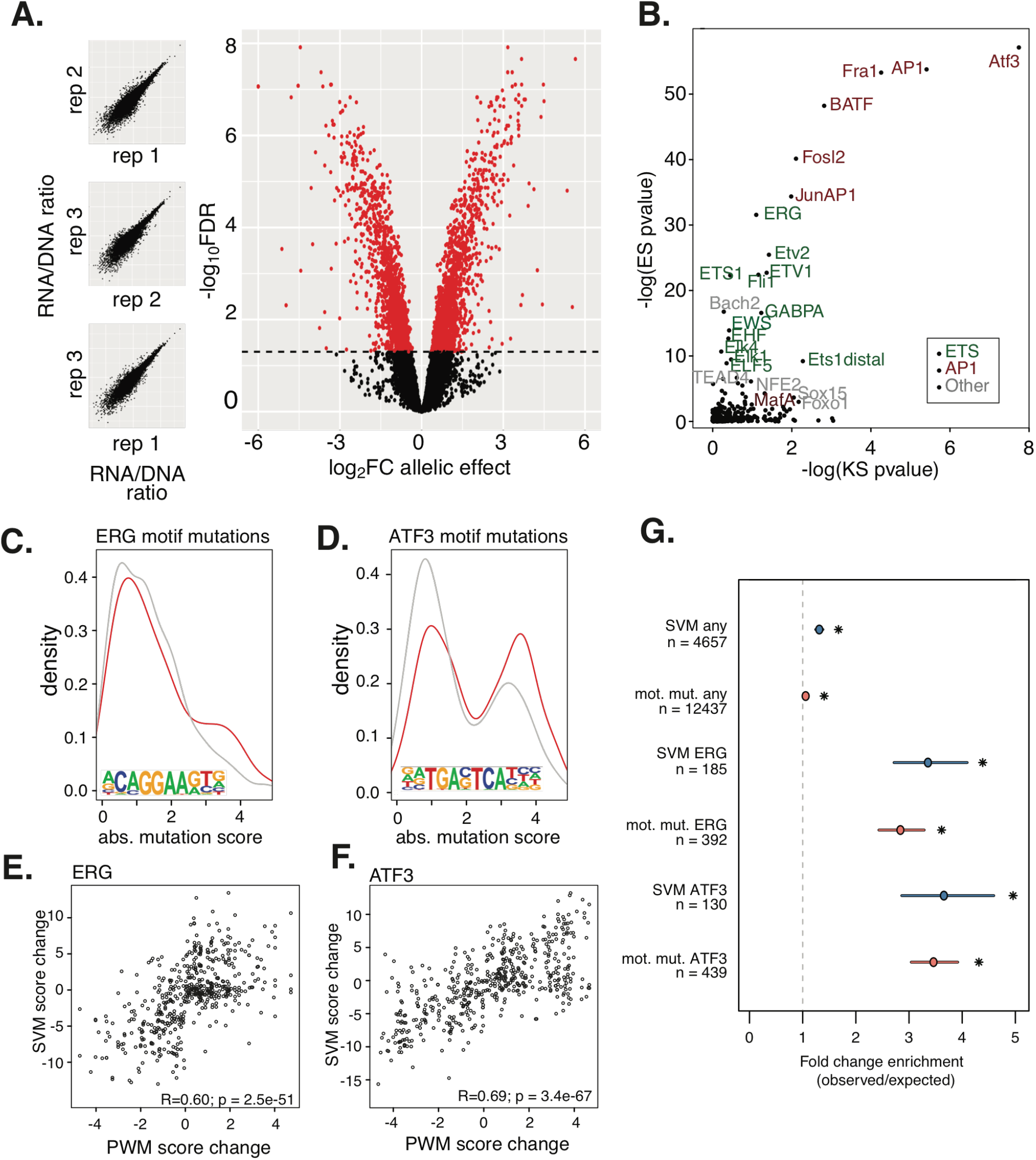
Allele specific enhancer function is associated with motif mutations. **A)** STARR-seq replicate RNA/DNA ratio correlations (left) and volcano plot of allele specific effects in STARR-seq with the log2 fold change on the x axis, and -log10 false discovery rate on the y-axis. **B)** Enrichment of transcription factor motif mutations compared to Kolmogorov– Smirnov testing between PWM motif mutation scores of validated vs non-validated SNPs. **C)** PWM motif mutation score density for ERG motif mutations in validated (red) and non-validated (grey). **D)** PWM motif mutation score density for ATF3 motif mutations in validated (red) and non-validated (grey). **E)** PWM motif mutation score (x-axis) versus the SVM score for ERG motif mutation SNPs. **F)** Motif mutation score (x-axis) versus the SVM score for ATF3 motif mutation SNPs. **G)** Enrichment of SVM (blue) and PWM motif mutations (orange) in the STARR-seq validated set.

It has been proposed that Support Vector Machine (SVM) models applied to *in vitro* TF-DNA binding measurements from SNP evaluation by systematic evolution of ligands by exponential enrichment (SNP-SELEX) experiments more accurately explain effects of allelic mutations to TF motifs than position weight matrices (PWM) for some TFs^19^. Therefore, we tested if SVM models were better indicators of allelic validation in our STARR-seq dataset than PWM mutation results. Among the 31k oligos detected, 29k had some predicted TF binding and among them 5,615 SNPs had SNP-dependent gain or loss of binding predicted. Besides ETS family factors, SVM-predictions were most successful for AP1- and CREB-like motifs (JDP2, ATF3, CREB1 etc; **Fig S3A)**. Importantly, we found significant correlation between SVM-based and PWM-based predicted allelic effects for many motifs, including ERG and ATF3 (**Fig. 2F-G, Fig. S3A**). Though predicted effects of both approaches were enriched for validated SNPs, SVM effects were slightly more associated with validation in STARR-seq. These data underscore the utility for both PWM and SVM methods for identification of functional SNPs in enhancers.

### Among epigenetic attributes, chromatin accessibility most significantly associates with allelic effects in STARR-seq

Epigenetic marks such as chromatin accessibility and histone modifications that are measured in native chromatin contexts are frequently used to signify genomic loci with enhancer function and to prioritize functional regulatory SNPs. Of SNPs that were included in our STARR-seq library, 5,155 had one of these indicators of regulatory function in epigenomes of HAECs, 1,418 of which validated in STARR-seq (**Fig. 3A**). Upon calculating enrichment scores, we initially found an enrichment of STARR-seq validated SNPs in HAEC genomic accessible regions, whereas SNPs within H3K27ac HAEC regions were minimally enriched (**Fig. 3B**). This led us to consider one major distinction between these data types; namely, the size of the peaks. ATAC-seq generates focal peaks (median 90 bps) whereas H3K27ac ChIP-seq produces distributed peaks (median 1492 bps) (**Fig. 3C**). We next hypothesized that functional SNPs are more likely to occur in H3K27ac regions when they are located near nucleosome-free regions (NFR) where TFs bind DNA. Restricting H3K27ac regions around their centers (100bp, 200bp, and 400bp) strongly improved enrichment scores for STARR-seq validated variants (**Fig. 3B**), demonstrating that SNPs closer to the center of the H3K27ac regions are more likely to perturb enhancer function than those further away. Given this result, used H3K27ac peaks restricted to 400 bp in subsequent analyses. Importantly, we found that SNPs in peaks having both ATAC-seq and H3K27ac modification were on average most enriched to validate in STARR-seq compared to either dataset alone (**Fig. 3B, bottom**).

**Figure 3.**
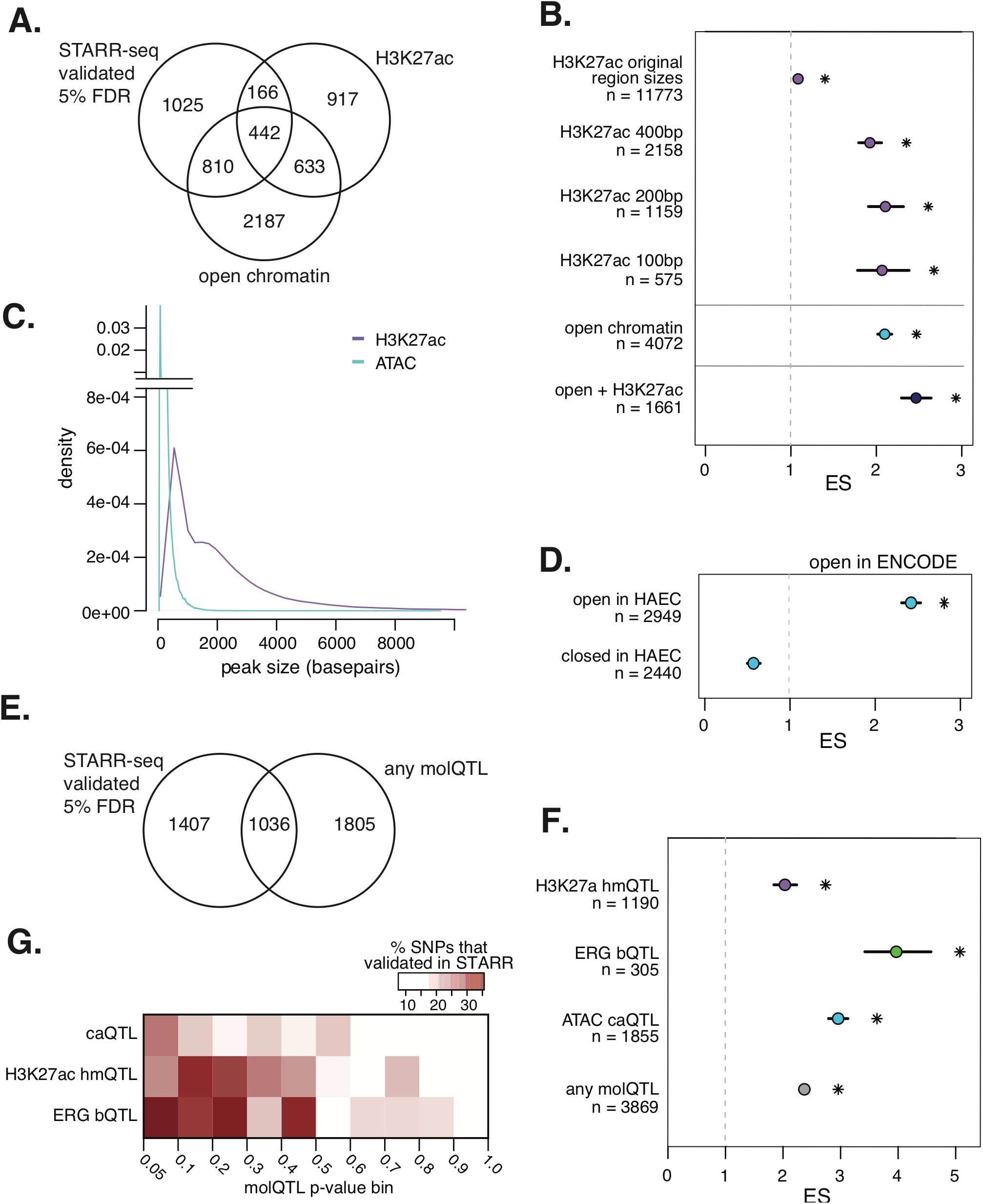
Epigenetic traits are linked to validation in STARR-seq. **A)** Venn-diagram comparing the number of SNPs in an open chromatin region in HAECs (bottom), in an H3K27ac peak (right), or validated in STARR-seq (left). **B)** Enrichment of SNPs in epigenetic trait peaks in the STARR-seq validated SNP set. H3K27ac (top 4) restricted to various sizes around the center of the peak. **C)** Peak size density of ATAC-seq peaks and H3K27ac ChIP-seq peaks. **D)** Enrichment of SNPs in ENCODE open regions that are shared and not shared with HAECs. **E)** Venn Diagram comparing the number of SNPs that are molQTLs (right) and validated in STARR-seq (left). **F)** Enrichment of molQTL SNPs in STARR-seq validated SNPs. **G)** Heatmap of number of STARR-seq validated SNPs in sub-significant molQTL bins.

We next asked whether SNPs in accessible regions in ECs were more likely to validate by STARR-seq than SNPs that were open in another cell type but not ECs. Using DNase I hypersensitivity peaks from 95 cell types in the Encyclopedia of DNA Elements (ENCODE)^20^, we found that open regions shared with HAECs are far more likely to validate than SNPs in open regions that are closed in HAECs (**Fig. 3D)**. Along these same lines, we observed diminished enrichment for HAEC-accessible SNPs when the same STARR-seq library was transfected into HepG2s^21^ (**Fig. S3**). These data are consistent with previous reports demonstrating that cell type is an important determinant for which regulatory elements, motifs, and allele-specific effects are detectable^11^.

### SNPs underlying molecular Quantitative Trait Loci frequently validate in STARR-seq

Molecular Quantitative Trait Loci (molQTLs) mapping is one approach to identify functional non-coding variants. In molQTL analysis, alleles/genotypes are tested for association with differences in epigenetic traits measured at a given locus. Significant molQTLs provide evidence for functional regulatory SNPs and often suggest a mechanism of action (e.g., an allele that perturbs TF binding). In prior work, we identified molQTLs for chromatin accessibility (caQTLs), H3K27ac histone modification (hmQTLs), and transcription factor ERG binding (bQTLs) in untreated HAECs. Of the SNPs input into our STARR-seq library, 2,841 were significant molQTLs (5% FDR), with 1,036 of these validating by STARR-seq (**Fig. 3E**). We found that each set of molQTLs were enriched for validation by STARR-seq (**Fig. 3F**), at magnitudes greater than epigenetic marks alone (**Fig. 3B**). Among molQTLs, caQTLs and ERG bQTLs were most enriched, supporting ERG’s importance in EC gene regulation. This result is replicated for bQTLs and caQTLs using Kolmogorov-Smirnov (KS) (p < 0.001; **Fig. S4**). Based on these analyses, we conclude that molQTLs effectively prioritize functional variants with focal epigenetic traits as most significantly associated. Still, we were interested that there is a subset of validated SNPs that are not molQTLs. We suspected this could be due to low statistical power, given the modest sample size (n∼20-50) for molQTL analysis. Consistent with this hypothesis, we observed an increased concentration of validated STARR-seq SNPs in p-value bins just above the molQTL significance threshold (**Fig. 3G**), leading us to conclude that sub-optimal molQTLs power in part explains the incomplete overlap between molQTLs and STARR-seq validated SNPs.

### Inflammatory Stimulus – GxE – IL-1b at 6 and 24 hours

Gene-by-Environment (GxE) interactions are one mechanism by which alleles can protect or predispose individuals to develop complex traits, including disease. In this study we modeled the effect of an inflammatory environment on endothelial cells in culture by pro-inflammatory cytokine IL1B stimulation, which is a known hallmark and driver of disease progression, including atherosclerosis^22^. Using the same STARR-seq library, we measured enhancer activity in teloHAECs exposed to IL1B for 6 and 24 hours, which loosely mimics early and late transcriptional responses to inflammation (**Fig. 4A**). In total, 3,297 regions had differential enhancer activity between treatments (2,695 upregulated by IL1B treatment, 602 downregulated by IL1B treatment) (**Fig. S3A**,**B**). As expected, the NF-kB motif was enriched in the regions with increased enhancer activity after IL1B treatment (**Fig. S3C**). Likewise, the ERG was highly enriched in regions with decreased enhancer activity after IL1B treatment (**Fig. S3C**). This is consistent with a previous report demonstrating that ERG protein is reduced upon pro-inflammatory stimulation^23^.

**Figure 4.**
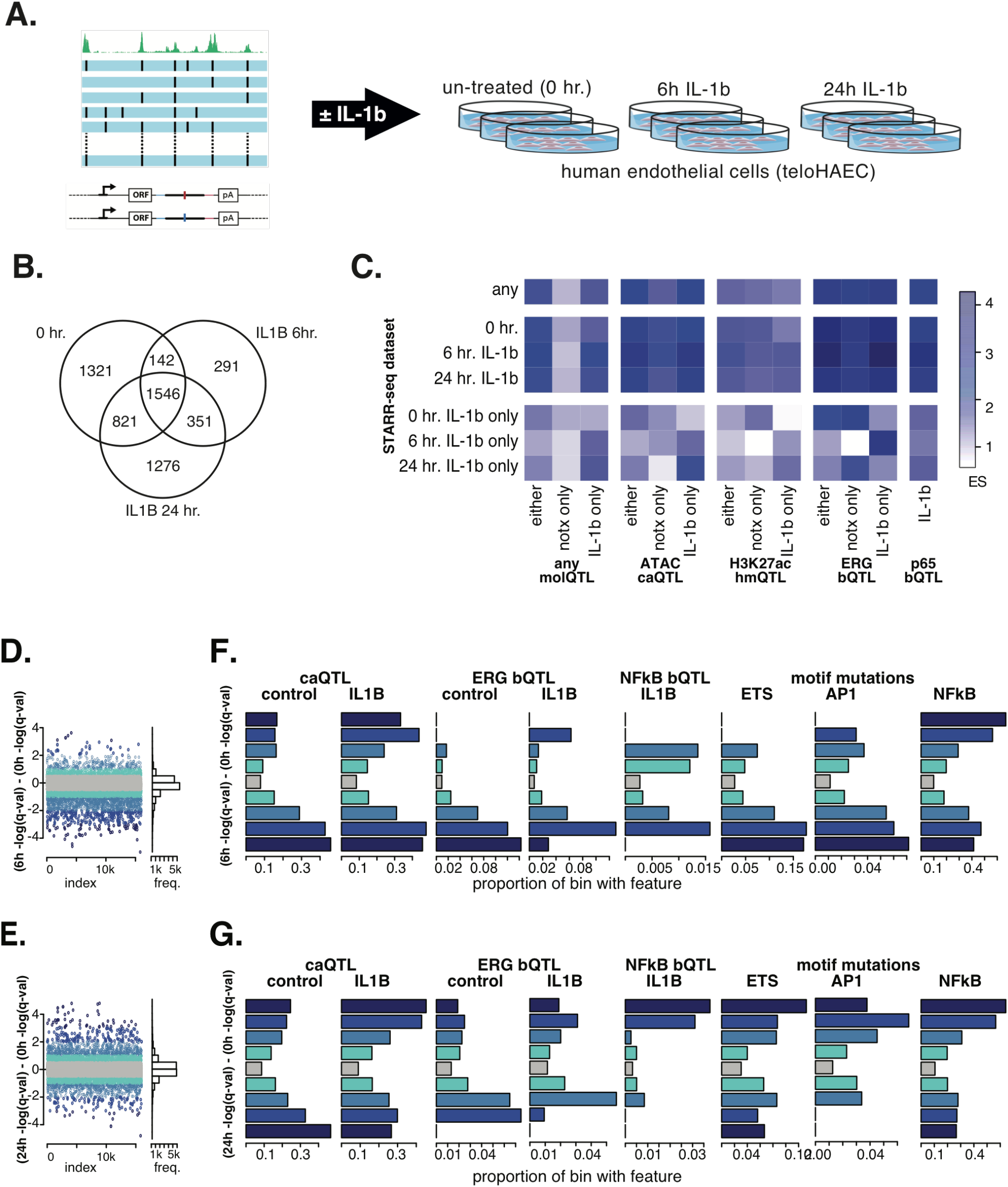
IL1B treatment trends with molQTLs and motif mutations. **A)** Diagram of treatment design. **B)** Venn Diagram of SNPs that validated in untreated (left), IL1B treated for 6 hours (right), and IL1B treated for 24 hours (bottom). **C)** Heatmap of enrichment scores comparing different treatment conditions between STARR-seq and the molQTLs. ‘Only’ categories refer to that treatments significant SNPs excluding any other treatments significant SNPs. **D)** Change in -log_10_(pvalue) between STARR-seq performed with 6 hour IL1B treatment verses untreated. Positive values are SNPs with more significance in 6h IL1B treatment. **E)** Change in - log_10_(pvalue) between STARR-seq performed with 24-hour IL1B treatment verses untreated. Positive values are SNPs with more significance in 24h IL1B treatment. **F)** Histograms of presence of molQTL SNPs and ERG/NF-kB/AP-1 motif mutations among bins along the y-axis of (D). **G)** Histograms of presence of molQTL SNPs and ERG/NF-kB/AP-1 motif mutations among bins along the y-axis of (E).

We identified SNPs with allele specific enhancer activity under IL1B treatment conditions and found 5,748 SNPs with significant effects in at least one treatment at 5% FDR. 1,546 SNPs (27% of those tested) had shared allelic effects across all treatment conditions (**Fig. 4B**). The 6-hour IL1B treatment had the least number of significant results (n = 2,330 SNPs), with the majority of those being similarly significant in either the 0-hour or 24-hour treatments. Thus, the 0-hour and 24-hour treatments uncover most SNPs that perturb enhancer activity.

We next evaluated relationships between allelic effects by STARR-seq (IL1B at 6 and 24 hrs) and molQTL results that were measured in HAECs after 4 hr IL1B treatment in HAECs^14^. Though there is enrichment in every STARR-seq dataset for every molQTL dataset tested (**Fig. 4C**), we observed patterns in the magnitudes with intriguing implications. For example, ERG bQTLs that are uniquely detected with IL1B treatment in HAECs are most enriched in the STARR-seq validated SNPs upon 6 hr IL1B treatment (**Fig. 4C**). This association is even more evident between IL1B-only ERG bQTLs and 6h IL1b-only STARR-seq SNPs (i.e., excluding SNPs significant in either 0hr or 24 hr IL1B treatment conditions; **Fig. 4C**). Additionally, H3K27ac hmQTLs that are uniquely detected in IL1B treatment are specifically enriched in STARR-seq validated SNPs after 24 hrs IL1B treatment (**Fig 4C**). This observation, that IL1B-restricted molQTLs tend to validate as IL1B-restricted STARR-seq signals with different kinetics lends insight into epigenetic mechanisms of action. Specifically, SNPs with effects at 6 hr IL1B are likely to be detected by perturbed TF binding (e.g., ERG), whereas SNP effects after 24 hr IL1B are evident from H3K27ac molQTLs.

In the enrichment analyses above, identification of SNPs altering enhancer activity relies on stark thresholding (e.g., at 5% FDR); further, identification of GxE regulatory SNPs requires comparing two thresholded datasets. We postulate this approach is not always biologically relevant. For example, SNPs near significance in one dataset, yet far from significance in another, likely exhibit GxE activity. We therefore chose to evaluate relationships between molQTLs and motif mutations using quantitative ratios of IL1B-treated and control (0h) STARR-seq summary statistics. In **Fig. 4D-E**, values near 0 on the y-axis indicate the SNPs having similar allelic effects between IL1B and 0h. SNPs with stronger effects in IL1B than 0h deviate toward larger y-values, and stronger effects in 0h than IL1B deviate toward smaller, negative y-values. Rightward-facing histograms in **Fig. 4D-E** demonstrate that most SNPs are near 0, and hence lack treatment-specific effects in STARR-seq. We considered regulatory GxE SNPs as those deviating from the 0-centered grey bin.

Interestingly, we found that SNPs with IL1B-specific STARR activity (positive on y) were more frequently caQTLs and ERG bQTLs in IL1B treatment conditions compared to their respective controls (**Fig. 4F**,**G)**. When comparing GxE SNPs by STARR at 6h IL1B vs 0h (**Figs. 4D**,**F**) to GxE SNPs at 24h IL1B vs 0h (**Fig. 4E**,**G**), we observed a trend that ERG bQTLs were most prevalent for GxE SNPs with stronger effects in 0h than with IL1B. Additionally, we found that frequencies of ETS motif mutations were pronounced in 0h relative to 6h STARR-seq SNPs, but the other way around for SNPs with 24h-specific perturbations relative to 0h. Lastly, we found that NFkB/p65 bQTLs, measured upon 4h IL1B treatment, were highly concentrated at GxE regulatory SNPs specific to 24h IL1B treatment (**Fig. 4G**). Correspondingly, the kB motif is most frequently mutated in the SNP set most specific to IL1B allelic activity. Taken together, these data demonstrate the prevalence of GxE on enhancer activity and underscore that SNPs affecting chromatin accessibility and binding of TFs can exert their effects only in particular cellular activation states.

### Validated functional SNPs fine-map GWAS signals for vascular diseases including abdominal aortic aneurysm and large artery stroke

To identify functional non-coding SNPs at GWAS loci, we cross-referenced our data generated in ECs with SNPs underlying GWAS signals for traits with appreciated vascular etiology. After restricting SNPs to those that validated by STARR-seq and had at least one significant molQTL, we report 89 high confidence functional regulatory SNPs that are associated with a variety of complex vascular diseases, 14 of which are associated with coronary artery disease (CAD) (**Table 1, Table S1**). One of these SNPs, rs17114036, has already been shown to alter enhancer function for the shear stress-induced transcript *PPAP2B*^24^ (Phosphatidate Phosphohydrolase Type 2b) (**Table 1**), thus serving as a positive control in our approach. We also see evidence for functional non-coding SNPs at the 9p21 CAD locus and the well-described *SMAD3* (Mothers Against Decapentaplegic Homolog 3) CAD locus. We find evidence that the *SMAD3* SNP rs17293632 affects enhancer activity in ECs, and further confirm that CRISPR-mediated deletion of the enhancer leads to significant reduction in the *SMAD3* expression in HAECs (**Fig. S6**); however, lack of an eQTL in HAECs and conflicting eQTL directions in GTEx obscures mechanistic interpretations. Two loci exhibited exceptional evidence for functional regulatory SNPs at enhancers that regulate target genes and modulate risk for: 1) Large Artery Stroke and Pulse Pressure, and 2) Abdominal Aortic Aneurysm. Respectively, we present evidence for functional non-coding SNPs, the *POU4F1 (POU Class 4 Homeobox 1)* locus, and rs13385499 and rs13382862 at the *LDAH* (Lipid Droplet Associated Hydrolase) gene locus.

**Table 1.** Selection of high confidence SNPs associated with vascular complex traits that are also a molecular QTL and significant in STARR-seq.

| rsID | Ref | Alt | Chrom | position (hg19) | eQTL genes (FDR<0.05) Both<br>HAEC Only<br>GTEx Only | Type molQTL<br>[.n = notx, .i =<br>i1b] (q-value,<br>higher tag allele) | Motif Mutated PWM | Predicted transcription factor binding disruption (SVM) | GWAS | STARR-seq pvals [$<0.05$ (Higher expression allele)]<br>0h<br>6h<br>24h |
| --- | --- | --- | --- | --- | --- | --- | --- | --- | --- | --- |
| rs6475604 | T | C | chr9 | 22052734 | CDKN2A\|CDKN2B | ATAC.n (0.02, T) |  | YY2 | Glaucoma\|CAD | 4.19E-02 (T)<br>0.227<br>0.234 |
| rs10757267 | G | C | chr9 | 22052810 | CDKN2B | p65.i (0.02, G) |  |  | Glaucoma\|CAD | 9.78E-05 (G)<br>3.45E-03 (G)<br>5.09E-04 (G) |
| rs17293632 | C | T | chr15 | 67442596 | SMAD3\|AAGAB\|PIAS1 | ATAC.n (2.5E-3, C) \| p65.i (4.2E-3, C) | Nkx6.1\|Lhx2\|ELF5\|EHF\|Lhx3\|Lhx1\|Fosl2\|JunAP1\|Fra1\|BATF\|AP1\|Atf3 | ATF3\|JDP2\|NFE2 | CAD\|Allergic rhinitis\|Asthma\|Atopic asthma\|Chronic inflammatory diseases ankylosing spondylitis Crohn's disease psoriasis primary sclerosing cholangitis ulcerative colitis pleiotropy\|Crohn's disease\|Eosinophil counts\|Hay fever and or eczema\|Inflammatory bowel disease\|Ulcerative colitis | 1.10E-04 (C)<br>9.66E-03 (C)<br>1.01E-03 (C) |
| rs13385499 | T | C | chr2 | 841284 | LDAH | p65.i (0.024, T) | CRX\|ZNF264\|Tlx\|Otx2\|GSC\|RUNX1 |  | Abdominal aortic aneurysm | 4.13E-05 (T)<br>8.57E-04 (T)<br>1.15E-04 (T) |
| rs13382862 | A | G | chr2 | 55576016 | LDAH | p65.i (0.029, A) | AP1\|BATF\|Atf3\|Fra1\|Fosl2\|Nkx3.1\|JunAP1\|Smad3\|Tbx5\|ERG | ERG\|FLI1 | Abdominal aortic aneurysm | 4.13E-05 (A)<br>8.57E-04 (A)<br>1.15E-04 (A) |
| rs4304924 | G | A | chr13 | 7846345 | POU4F1<br>RNF219-AS1\|C13orf7<br>RNF219\|RP11-52L5.6 | H3K27ac.n (3.4E-8, A)\|H3K27ac.i (0.011, A) | CRX\|GSC\|GATA3\|Gata4 |  | Large artery stroke\|Pulse pressure | 1.54E-03 (A)<br>0.125<br>2.91E-02 (A) |
| rs17114036 | A | G | chr1 | 56962821 | PPAP2B | p65.i (0.03, G) |  |  | Coronary artery disease\|Coronary artery disease or ischemic stroke\|Coronary artery disease or large artery stroke\|Coronary heart disease | 1.69E-02 (G)<br>3.68E-03 (G)<br>3.49E-03 (G) |

On chromosome 13, at a GWAS signal for Large Artery Stroke and Pulse Pressure^25; 26^, we identified that alleles of rs4304924 associate with differential regulatory activity and allele specific expression of *POU4F1* located ∼61 kB upstream (**Fig. 5A**). *POU4F1* is a gene that encodes for a homeobox transcription factor with described roles in neuronal differentiation^27^ and yet unknown roles in vasculature. We observed differential regulatory activity as an H3K27ac hmQTL and allele-specific expression in 0hr IL1B STARR-seq (**Fig. 6B**,**C, Table 1**). The rs4304924 reference allele (G) coincides with homeobox motif mutations, among others (**Table 1; Fig. 5D**). We also observed an eQTL for *POU4F1* and rs4304924 in HAECs, in addition to significant *POU4F1* eQTLs in Coronary, Tibial, and Aortic Artery GTEx tissues, thereby confirming vascular effects (**Fig. 6E**,**F**). Taken together, our data prioritizes further inquiry into what role *POU4F1* plays in the arterial vasculature.

**Figure 5.**
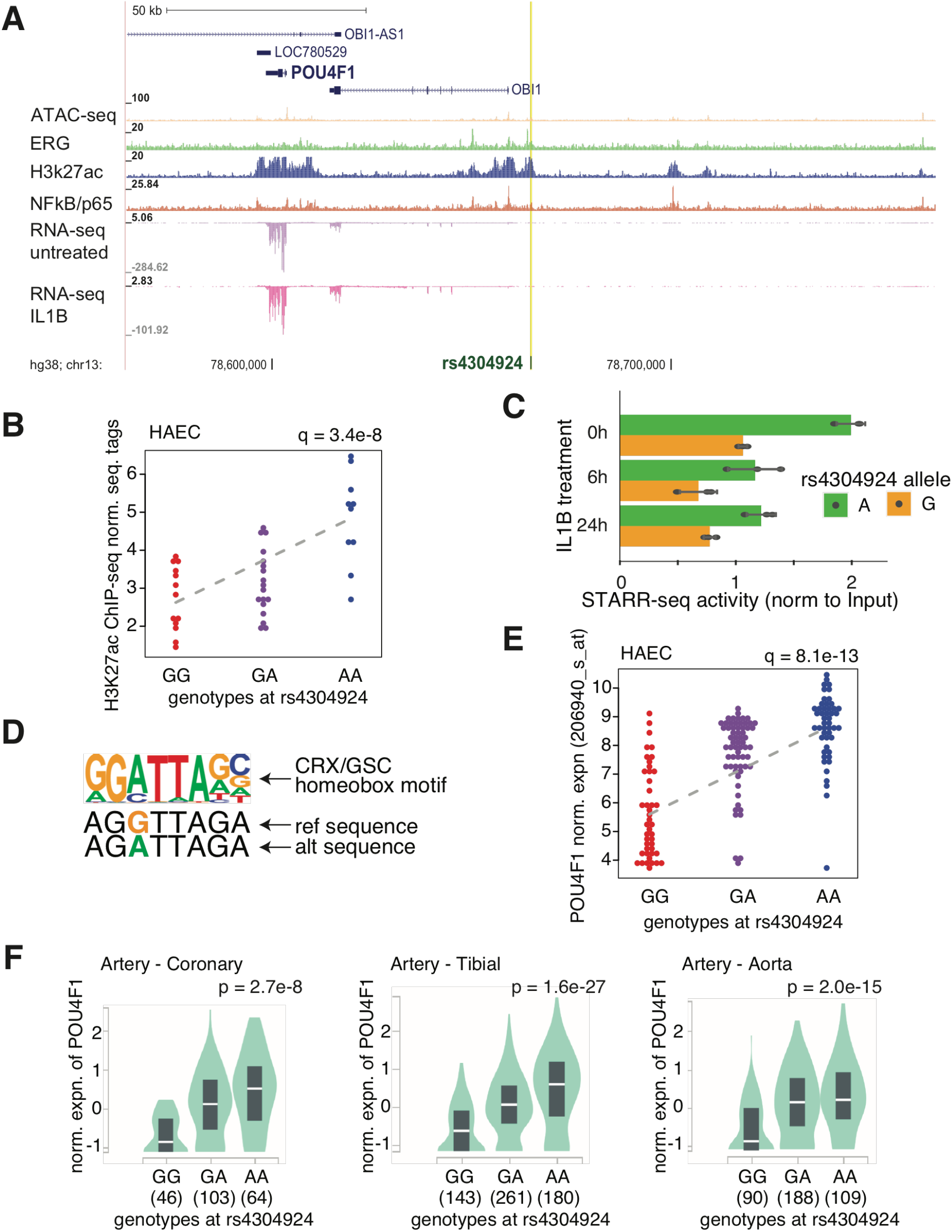
Large artery stroke and pulse pressure associated locus at gene POU4F1. **A)** Genomic region containing the putative enhancer where rs4304924 (yellow) resides and the surrounding genes. Tracks below display H3K27ac, ATAC-seq, ERG binding, p65 binding, and RNA-seq expression from HAECs. **B)** H3K27ac tags in HAECs compared between genotypes at rs4304924 (RPKM normalized). **C)** STARR-seq allele specific expression for rs4304924 across the different treatment points. **D)** PWM for the CRX/GSC motif and the motif sequences created by the two alleles at rs4304924 **E)** POU4F1 gene expression in HAECs by microarray compared between genotypes at rs4304924 **F)** POU4F1 gene expression by genotypes at rs4304924 within three arterial tissues from GTEx.

**Figure 6.**
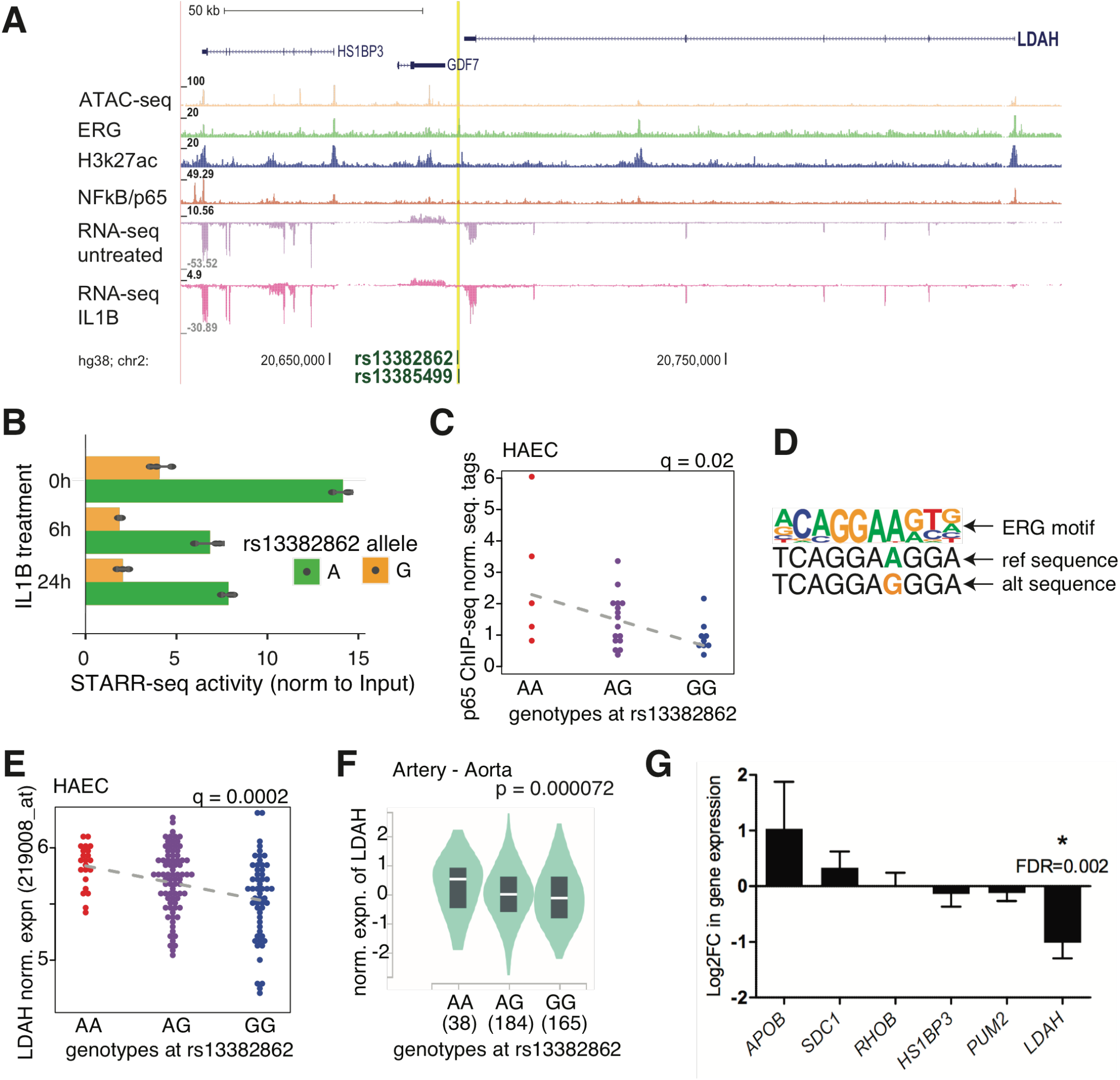
Abdominal aortic aneurysm associated locus at gene LDAH. **A)** Genomic region containing the putative enhancer where rs13382862 and rs13385499 (yellow) reside and the surrounding genes. Tracks below display H3K27ac, ATAC-seq, ERG binding, p65 binding, and RNA-seq expression from HAECs. **B)** STARR-seq allele specific expression for rs13382862 across the different treatment points. **C)** p65 (NF-kB) tags in HAECs compared between genotypes at rs13382862 (RPKM normalized). **D)** PWM for the ERG motif and the motif sequences created by the two alleles at rs13382862 **E)** LDAH gene expression in HAECs by microarray compared between genotypes at rs13382862 **F)** LDAH gene expression by genotypes at rs13382862 within three arterial tissues from GTEx **G)** Effect of rs13385499 and rs13382862 carrying enhancer deletion on the expression of genes within 1 Mb. Only *LDAH* demonstrates significant repression. DESeq2 was used to compare the enhancer deleted samples (n=4) to the controls (n=8). The data is represented as log2 fold change and standard error for the estimated coefficients on the log2 scale.

At a GWAS locus for Abdominal Aortic Aneurysm^28^, two SNPs, rs13385499 and rs13382862, validated in STARR-seq at all time points, and underlie p65 bQTLs in HAECs (**Fig. 6A-C, Table 1**). SNPs rs13385499 and rs13382862 are in LD with each other (EUR R^2^ = 0.79; HAEC population R2=0.89)^29^. Both SNPs cause multiple motif mutations, including for RUNX (rs13385499) and ERG (rs13382862) motifs, which directionally coincide with the p65 bQTL (**Fig. 6D**). These SNPs are at the 3’ end of the *LDAH* gene, and demonstrate association with transcript expression levels in HAECs (**Fig. 6A**,**E**). In addition, both SNPs are eQTLs for *LDAH* in the same direction in multiple GTEx tissues including for Artery Aorta (rs13382862) (**Fig. 6F**). *LDAH* has been associated with cholesterol deposition in atherosclerotic plaques^30^; however, the role of *LDAH* in ECs has not been well described. Unlike *POU4F1, LDAH* is highly expressed in teloHAECs, and targeted deletion of the 456 bp enhancer (**Table S2; Fig. S7**) carrying both SNPs resulted in decreased *LDAH* expression with minimal effect on other nearby genes (**Fig. 6G**). Together, these data underscore that rs13385499 and rs13382862 are credible functional SNPs for Abdominal Aortic Aneurysm and prescribe a role for *LDAH* in the disease process.

## Discussion

In this study, paired MPRA and epigenetic data from genetically diverse HAECs enabled a series of discoveries. Importantly, it allowed us to specify which genomic features most often describe SNPs with functional effects on enhancer activity. Our findings, in **Fig. 2** and **3**, suggest that functional non-coding SNPs likely have one or more of the following attributes: 1) an allelic mutation to a TF binding motif for a factor with an important role in the cell type and/or cell state of interest, 2) genomic location in a region marked by accessible chromatin and H3K27ac, 3) association to an epigenetic measurement in *cis-* (molQTL), with effects on chromatin accessibility and TF binding being most significant. Using IL1B as a modulator of cell state and environmental simulant, we observe robust evidence for context-specific regulatory SNP effects, thereby underscoring the prevalence of GxE effects on enhancer function. Lastly, we fine-map functional non-coding SNPs at GWAS loci for vascular diseases through integration of molQTL mapping, STARR-seq, and eQTLs from HAECs and GTEx tissues. These findings are discussed below.

We find that AP-1 ad ETS motif mutations are the most enriched at functional regulatory loci detected in our STARR-seq experiment (**Fig. 2**). Previous findings indicate that perturbed ERG and JUN bindings are likely affected by these motif mutations, although other members of the ETS and AP1 families are almost certainly affected as well, as many members are highly expressed in ECs^5^. The enrichment of functional mutations in ETS and AP1 motifs is consistent with other reports demonstrating that functional mutations in regulatory elements often reveal motifs of TFs that are selectively active in the respective cell types^13^. Additionally, both AP-1 and ETS factors are demonstrated to be pioneering factors, able to establish accessible chromatin, and ‘select’ primed and active enhancers to which collaborating TFs can bind. This was shown previously in ECs and macrophages; namely, that genetic variants that perturb the pioneering binding function of AP-1 and ETS factors consequently affect binding of collaborating and signal-dependent TFs^5; 31; 32^. Irrespective of cell type, AP-1 motifs have been shown to produce high enhancer activities^33^, whereas ETS factors seem to show high episomal activity^34^ suggesting that these motifs might be more evident in validated elements by MPRAs. Still, based on similar enrichment values of STARR-validated SNPs, we show that SVM-based ERG motif mutations have slightly lower enrichment among validated SNPs than ERG bQTLs. This demonstrates the potential of improved TF motif models^19^ to identify functional variants in the future.

Still, many SNPs that fit the epigenetic and/or mutation criteria outlined above did not validate in STARR-seq. For example, considerable numbers of SNPs in accessible and/or H3K27ac-marked chromatin that did not validate by STARR-seq (**Fig. 3A**). Perhaps more confusing is that the majority (64%) of SNPs with molQTLs (i.e., prior evidence of allelic effects) in HAECs did not validate in STARR-seq (**Fig. 3E**). Possible explanations include that: (1) causal alleles of weak effect may fall below limit of detection in STARR-Seq, (2) detection requires longer sequence context not captured on the 198 bp oligo. Also, (3) SNP effects might require the endogenous genomic position, chromatinization and sequence context not captured by episomal assays and, (4) silencing effects of alleles are likely undetectable due to the low basal activity of the minimal promoter used. These are limitations that should be addressed in future experimental iterations. Further insight into such discrepancies between genomic features and MPRA results will be necessary to accelerate discovery into the functional non-coding genome.

Genetic differences between individuals can profoundly alter how cells respond to environmental cues. By conducting STARR-seq at three IL1B treatment timepoints in teloHAECs (0h, 6h, 24h), we were able to observe the striking relationship between cell activation state and enhancer SNPs with allelic effects (**Fig. 4**). Upon comparison to molQTLs measured in untreated and IL1B treatment (4h), we see that SNPs at IL1B-specific molQTLs are enriched in IL1B-specific validated STARR-seq SNPs with higher proportions of NFkB motif mutations in the IL1B-specific SNPs (**Fig. 4D**). Together, these findings confirm that the genomic consequence (molQTLs) of many SNPs is replicated episomally (STARR-seq) and that the effect of non-coding SNPs are often cell state-specific. In addition, our results illustrate how pre-existing genetic effects on chromatin could propagate to enhancer activity. To this end, we observed that ERG bQTLs were most prevalent for GxE SNPs that demonstrated stronger allele specific enhancer activity in unstimulated conditions. This supports the observations that large fraction of stimulus-specific eQTLs perturb enhancer priming by lineage determining transcription factors^31; 32^. In contrast, we found that frequencies of ETS motif mutations were more pronounced in 0h relative to 6h STARR-seq SNPs, but the opposite for 24h-stimulation. This could result from another ETS family member than ERG binding the motif under IL1B signaling or reflect the changes in ERG expression upon inflammatory stimulus^23^. Alternatively, this could be explained by different kinetics of epigenetic changes and STARR-Seq transcription or by differences in early and late response to inflammatory stimulus. For example, the allele specific enhancer activity at 24 hr IL1B was more evident from H3K27ac molQTLs, whereas the 6h effects were more likely to be detected by perturbed TF binding. Whether these correlations change if the TF binding or epigenetic changes are measured at later timepoints warrant future research. The implication of our findings is that a comprehensive understanding of non-coding functional SNPs will require experimental observations from comprehensive sets of cell types and cell states. Furthermore, models of gene regulation will need to consider functions of all relevant enhancers, cell activation state, as well as the genetic variants at the relevant enhancers. Such context specificity exemplifies the vast complexity of gene regulatory networks.

A major motivation for this study was to fine-map functional non-coding variants at disease loci for complex human vascular diseases. **Table 1** is our credible functional SNP list with corresponding lines of evidence for each SNP that meets genome-wide significance for a disease trait with vascular relevance. There are SNPs present in the dataset that have a molQTL, and validated in STARR-seq, yet do not have any documented eQTL in HAECs or GTEx. These SNPs could potentially operate through mechanisms not detected by poly-A-selected mRNA transcripts (i.e., splicing, regulation of non-polyadenylated RNA transcripts). Or perhaps, we indeed measured functional SNP effects, yet the SNPs reside in pseudo-enhancers that do not directly target any gene. Furthermore, we find it interesting that some SNPs are in our credible set have a GTEx eQTL but not an HAEC eQTL. This set potentially represents a group of SNPs for which we lack the power to detect eQTLs, or they could be eQTLs in another cell-type. For example, at 9p21, we observe two validated SNPs, rs6475604 and rs10757267, with molQTLs. Whereas these SNPs are not associated to any transcripts in HAECs, they are each eQTLs for *CDKN2B* in skeletal muscle in GTEx (**Fig. S6A-D)**. We also identified rs17293632 that is an AP1 motif mutation and has been previously identified as candidate causal variant for *SMAD3* in smooth muscle cells^35-38^. Despite absence of cis-eQTL to *SMAD3* in HAECs, we demonstrate that deletion of the rs17293632-carrying enhancer does abrogate *SMAD3* expression in ECs (**Fig. S6I; Fig. S7**). Interestingly, GTEx shows significant associations to *SMAD3* in thyroid and esophagus mucosa in opposing directions, indicating that AP1 motif binding factors could recruit either repressive or activating complexes (**Fig. S6G**,**H**). This could mean that the same variant acts through many cell types to regulate the process of atherosclerosis possibly through transforming growth factor β (TGF-β) signaling^39^. The current study is the first study to implicate this SNP in ECs. Still, further investigation will be required to dissect mechanisms of action at this interesting locus.

One of the most interesting findings from our study was the in-depth characterization of a locus associated to Large Artery Stroke and Pulse Pressure. We present strong evidence for the functional role of rs4304924 in an element regulating expression of *POU4F1* (**Fig. 5**). The G allele of this SNP mutates a homeobox TF motif, exhibits diminished H3K27ac in the adjacent chromatin, and has reduced *POU4F1* expression compared to the A allele. This eQTL is replicated in all three arterial GTEx tissues. To our knowledge, there are no publications linking this transcript with function in the arterial vasculature. The other most interesting finding is at a locus associated to Abdominal Aortic Aneurysm where we find two SNPs, rs13385499 and rs13382862, whose alternate alleles correspond to diminished p65 binding and expression of the nearby gene *LDAH* (**Fig. 6**). The alternate alleles mutate TF motifs, and rs13382862 is likewise an eQTL for *LDAH* in GTEx’s Artery Aorta dataset. The two SNPs are in high LD in our HAEC population, although in the greater European population they are in moderate LD (R2=0.8). Given that rs13382862 causes an ETS motif mutation and is an eQTL for *LDAH* in GTEx and HAECs, we posit that this SNP has a greater functional consequence. *LDAH* itself has been mostly described in macrophages where upregulation of *LDAH* is linked with a reduction in intracellular cholesterol^30; 40^. Additionally, it has been shown to be highly expressed in macrophage laden atherosclerotic plaques^30^. In this study, we present a mechanistic link between inherited non-coding variation, *LDAH* expression, and risk for Abdominal Aortic Aneurysm.

Taken together, the implication of our findings is that a comprehensive understanding of non-coding functional SNPs will require experimental observations from comprehensive sets of cell types and cell states. Such context specificity exemplifies the vast complexity of gene regulatory networks.

## Methods

### Generating Haplotypes for STARR library

The following computational pipeline was used to generate 198 bp sequences representing up to 5 haplotypes at each locus of interest for including in the STARR-seq library. First, genomic loci of interest were identified using multiple criteria including overlap of HAEC epigenetic features and sequence analysis and formatted as HOMER peak files^41^. Second, phased alleles along haplotypes, in VCF format, were utilized from our previous study of 53 genotyped, imputed, and phased individuals that generated the HAEC epigenetic data^14^. Note that alleles were only included if they had Minor Allele Frequencies greater than 5% in this population. Third, the HOMER sub-program *annotatePeaks*.*pl* was used by inputting the peak file from step 1 along with the options “-vcf phased.vcf.file -size given” which outputs another HOMER formatted peakfile with columns noting the bp positions within each peak and alleles of each haplotype. Fourth, the sequence of the reference hg19 human genome was retrieved within each peak boundaries using the R package *seqinr()*^42^. A custom R script was then used to iterate through each peak to paste custom sequences together for each haplotype. Specifically, strings of non-polymorphic sequence were separated from polymorphic alleles using coordinates in the previous peak file, and then these were pasted together again for each haplotype. Fifth, resulting sequences were compared along each haplotype, and duplicated sequences were removed, which sometimes arose when peak sequences were identical between haplotypes. Custom code is available upon request.

### Massively parallel reporter assay

SNPs with functional non-coding evidence shown in **Fig. S1** were included in the library. In addition, 100 scrambled regions and 100 random negative regions that do not overlap any chromatin marks in the ENCODE database^20^, were selected. The following computational pipeline was used to generate 198 bp sequences representing up to 5 haplotypes at each locus of interest for including in the STARR-seq library. First, genomic loci of interest were identified using multiple criteria including overlap of HAEC epigenetic features and sequence analysis and formatted as HOMER peak files^41^. Second, phased alleles along haplotypes, in VCF format, were utilized from our previous study of 53 genotyped, imputed, and phased individuals that generated the HAEC epigenetic data^14^. Note that alleles were only included if they had Minor Allele Frequencies greater than 5% in this population. Third, the HOMER sub-program annotatePeaks.pl was used by inputting the peak file from step 1 along with the options “-vcf phased.vcf.file -size given” which outputs another HOMER formatted peakfile with columns noting the bp positions within each peak and alleles of each haplotype. Fourth, the sequence of the reference hg19 human genome was retrieved within each peak boundaries using the R package seqinr()^42^. A custom R script was then used to iterate through each peak to paste custom sequences together for each haplotype. Specifically, strings of non-polymorphic sequence were separated from polymorphic alleles using coordinates in the previous peak file, and then these were pasted together again for each haplotype. Fifth, resulting sequences were compared along each haplotype, and duplicated sequences were removed, which sometimes arose when peak sequences were identical between haplotypes. Custom code is available upon request.

The hSTARR-seq_ORI plasmid (Addgene, #99296)^43^ was used as a backbone for the plasmid constructs. DNA inserts, 230 bp long and containing 198 bp of enhancer variant sequence, were synthesized by Agilent. The oligos were designed to have a 2 bp barcode in the 5’ end of the enhancer sequence and 15 bp matching to Illumina NGS sequencing primers in both ends. First round of emulsion PCR using Micellula DNA Emulsion & Purification Kit (Roboklon) was performed to complete the sequencing primers and to double-strand the oligos. The second round was used to amplify the material. The plasmid was linearized using AgeI and SalI restriction enzymes and inserts were cloned to the linearized plasmid in 17 reactions using the standard InFusion cloning (Clontech) protocol. The cloned DNA library was transformed to XL-10 gold ultracompetent bacteria (Agilent) in 15 reactions. Plasmid was purified using EndoFree Maxiprep kit (Qiagen). Immortalized human aortic endothelial cells (teloHAECs) were purchased from ATCC and cultured in Vascular Cell Basal Medium (ATCC PCS-100-030), supplemented with Vascular Endothelial Cell Growth Kit-VEGF (ATCC PCS-100-041), 100 U/ml penicillin and 100 µg/ml streptomycin. Cells were incubated at 37°C in 5% CO2 and passaged every 3 days until the number of cells needed for the experiment was achieved.

The plasmid library was transfected in triplicates to 90 million cells/replicate using Neon transfection system (Invitrogen). The cells were detached with trypsin, centrifuged, washed with PBS and resuspended to resuspension buffer R at 5×10^7^ cells/ml. The cell suspension was mixed with 15 µg of STARR-seq library/reaction and subjected to electroporation using the 100 µl tip and three 1350V pulses of 10 ms width. The cells were allowed to recover from electroporation for 24 hours, after which they were treated with inflammatory stimulus (IL1β, 10 ng/ml). The cells were divided to 3 treatment groups, each with 30 million cells: 6 h stimulus, 24 stimulus and non-stimulated control. Cells were harvested 48 h post-transfection and RNA was extracted using RNeasy midi kit (Qiagen). Messenger-RNA was purified from bulk RNA using Dynabeads Oligo(dT)25 beads (Invitrogen) with 2:1 bead to total RNA volume ratio. The purified mRNA was treated with Turbo DNaseI (Ambion) and purified using RNeasy MinElute clean up kit (Qiagen). Reverse transcription was performed using UMI-primers. Unique molecular identifiers (UMIs) were added during cDNA synthesis to tag identifiable replicates of the constructs, which improves the data analysis by accounting for PCR duplicates^44^. The samples were pooled, RNase A -treated and the cDNA was purified with AMPure XP beads using a 1.8:1 beads to cDNA ratio. The libraries were amplified using junction PCR. The junction PCR for RNA library was performed with junction_RNA_fwd and junction_RNA_rev primers^43^, which allow amplification of correctly inserted enhancer sequence cDNA. The jPCR products were purified using AMPure XPbeads with beads to sample ratio of 0.8. A second PCR step was run to add the index primers for sequencing (NEBNext Multiplex Oligos for Illumina Dual Index Primers Set 1 and 2). PCR products were purified using SPRIselect beads (Beckman) (bead to sample ratio 0.8). Next generation sequencing was performed on Illumina NextSeq 500 platform as paired end 75 cycle dual index runs using the following parameters: Read1: 37bp (insert sequence), Index1: 10bp (contains the UMI instead of an i7 index), Index2: 8bp (i5 index for demultiplexing), Read2: 37bp (insert sequence). Raw data from STARR-seq experiments is available at NCBI GEO (accession GSE180846).

Sequencing read demultiplexing was carried out using the i5 index (Index2) only, and the Index1 read (containing the UMI) was extracted as regular sequence read. UMI-tools version 1.0.1^45^ was used to remove any reads where the UMI did not match the expected sequence pattern RDHBVDHBVD52. The remaining reads with valid UMIs were aligned to a custom reference genome consisting of all the oligonucleotide sequences included in the library. Alignment was performed with the STAR aligner by running the nf-core RNA-Seq pipeline version 1.4.2^46^. UMI deduplication was performed to the resulting BAM files using the UMI-tools default method. Reads mapping uniquely and strand-specifically to a library oligonucleotide were summed using featureCounts (Subread Python package version 1.6.5)^47^. To identify enhancers displaying allele-specific expression, STARR-seq read counts for each variable position were summed by variant allele (reference or alternative) and the mpralm R package version 1.4.1^18^ was used.

### PWM-based and SVM-based TF motif mutation analysis

To test for evidence of functional SNPs by identifying alleles distinguishing DNA motifs, we used the MMARGE software package^48^. The human reference hg19 build^49^ was input with genotyped and imputed SNPs of our HAEC population (dbGAP Study Accession: phs002057.v1.p1) to create reference builds in MMARGE for reference and alternate allele MMARGE’s the ‘prepare_files’ function. A MMARGE-formatted peakfile was created using reference genomic chromosome and position boundaries for all oligos input into the STARR-seq library. These were input into MMARGE’s mutation_analysis function, using the ‘-keep’ flag to save temporary output files, along with all TF motifs available in the HOMER^41^ database. The temporary files returned were queried to create a unique list of genomic positions, motifs detected, and motif scores for the reference and alternative genome sequence builds. The positions for the alternate genome were shifted to reference coordinates to remove positional differences resulting from indels between reference and alternate builds. This was achieved with tabix^50^ by adding back the difference in allele base pair lengths between reference and alternate alleles to the alternate positions. Motif scores were compared for corresponding motif positions and the difference is reported as the delta PWM in this study. When a motif was only detected in either the reference or alternate, the minimum PWM detection score (from HOMER) was used for the absent genome. As an alternative approach for the prediction of TF binding disruption due to genetic variants, recently published deltaSVM models^19^ based on *in vitro* protein-DNA binding data were run for all variants in the current study. All 94 high-confidence TF binding models published by the original authors were included and run with the authors’ recommended thresholds for sequence binding and allelic disruption.

### Enrichment testing

To calculate enrichment of any dataset in the STARR-seq significant datasets, we counted how many SNPs were both STARR-seq positive, and positive for the test dataset. This number became the experimental value (observed). We then approximated the expected number of SNPs that would satisfy these criteria due to the selective nature of the library creation. This value roughly equals the proportion of SNPs in the library that are in the test dataset multiplied by the total number of SNPs that came out STARR-seq positive (num_pos). This value became our expected value (expected). The observed was compared to expected using a one-sided binomial test in R as follows: binom.test(x = observed, n = num_pos, p = (expected/num_pos), alternative = “greater”). We then calculated an adjusted p-value using Benjamini-Hochberg correction and any dataset that reached an FDR threshold of 0.05 was considered significantly enriched in the STARR-seq dataset in question.

### ENCODE comparison

To compare enrichment between HAEC open regions verses regions that are not open in HAECs but are open in other cell types, the ENCODE track was downloaded in bed file format (encode.bed) from the UCSC genome browser^51^. This file was processed using HOMER^41^ to restrict peak sizes to 200 bp using command (annotatePeaks.pl encode.bed hg19 -size 200). This peak file (encode.peaks) was then separated into those that are shared with HAECs based on our ATAC-seq data across 44 individuals (atac.peaks) also using HOMER as follows (mergePeaks -cobound 1 encode.peaks atac.peaks).

### Kolmogorov-Smirnov Testing

As a validation of the enrichment testing, we verified the associations by using Kolmogorov-Smirnov testing between the QTL analysis FDRs of the STARR-seq validated sets of SNPs vs the SNPs that did not validate. The FDRs for one dataset (eg. ERG binding QTL FDRs) were pulled for all of the SNPs that validated (valid) and all SNPs that did not validate (noValid). The R code to perform testing is as follows using command from package ‘stats’: ks.test(valid, noValid, alternative = “less”). Plotting of the densities for graphical representation of the distributions tested were done by using the ‘density’ command from package ‘stats’. Cumulative distributions were created with command ‘Freq’ from package ‘DescTools’.

### CRISPR validation experiments

CRISPR/Cas9-mediated deletion of target regions was performed using the Alt-R CRISPR-Cas9 System (Integrated DNA technologies; IDT, Coralville, Iowa). Briefly, CRISPR/Cas9 single gRNAs (**Table S2)** flanking the target enhancers were designed using an online CRISPR design tool (https://eu.idtdna.com/site/order/designtool/index/CRISPR_SEQUENCE) and ordered from IDT as crRNAs. These crRNAs were annealed to a tracrRNA (#1072532, IDT) and complexed with Cas9 endonuclease (S.p. HiFi Cas9 Nuclease V3, #1081060, IDT) to form the ribonucleoprotein complex (RNP). The RNP complexes were then delivered into 150 000 teloHAEC (ATCC) cells per replicate by electroporation using the Neon transfection system (Invitrogen) with a 1350 V pulse of 30 ms width, following IDT CRISPR genome editing protocol for RNP electroporation, Neon Transfection system. Two days later, transfected cells were collected for RNA and gDNA. In order to analyze the efficiency of enhancer deletion, genomic DNA was extracted using NucleoSpin tissue kit (Macherey-Nagel, Düren, Germany) and amplified by PCR using specific primers (**Table S2**) flanking the deletion sites and DreamTaq DNA Polymerase; EP0701,Thermo Fisher Scientific). PCR products were then analyzed by electrophoresis in 1% agarose gel. To analyze the effects of enhancer deletion on cis target gene expression, the RNA was extracted using RNeasyPlus Micro kit ;74034, QIAGEN) RNA library preparation was carried out using QuantSeq 3’ mRNA-Seq Library Prep Kit FWD for Illumina (Lexogen, Vienna, Austria) according to the manufacturer’s instructions. The resultant library was quantified using Qubit dsDNA HS Assay Kit (Q32854,ThermoFisher Scientific) and its quality was checked with Bioanalyzer using High Sensitivity DNA Kit (5067-4626, Agilent Technologies). Individual libraries were pooled in equimolar ratio (4 nM for each) and sequenced with the Nextseq 550 platform (Illumina, San Diego, California) in a single end 75 cycles high output run. The effect of CRISPR deletion on gene expression was studied for all the cis candidate genes within 1 Mb of the deleted region. The positive and negative guide RNA transfected cells were used as controls (n=4 each). DESeq2^52^ was used for the statistical analysis and FDR<5% was considered significant.

## Data Access

Raw data for STARR-seq experiments is available at NCBI GEO accession GSE180846. Reviewers can access the records by entering the token: yvyxqcywnnktrgt.

## Acknowledgements

This research was supported by the National Institutes of Health (NIH) (R01HL147187 to C.E.R. and T32 HL007249-42 to L.K.S)., the European Research Council (ERC) under the European Union’s Horizon 2020 research and innovation program (Grant No. 802825 to M.U.K), the American Heart Association (20PRE35200195 to L.K.S.), the Academy of Finland (Grants Nos. 287478 and 319324 to M.U.K.), the Finnish Foundation for Cardiovascular Research, the Sigrid Juselius Foundation, and the Doctoral Program of Molecular Medicine at University of Eastern Finland.

---

**Figure S1.**
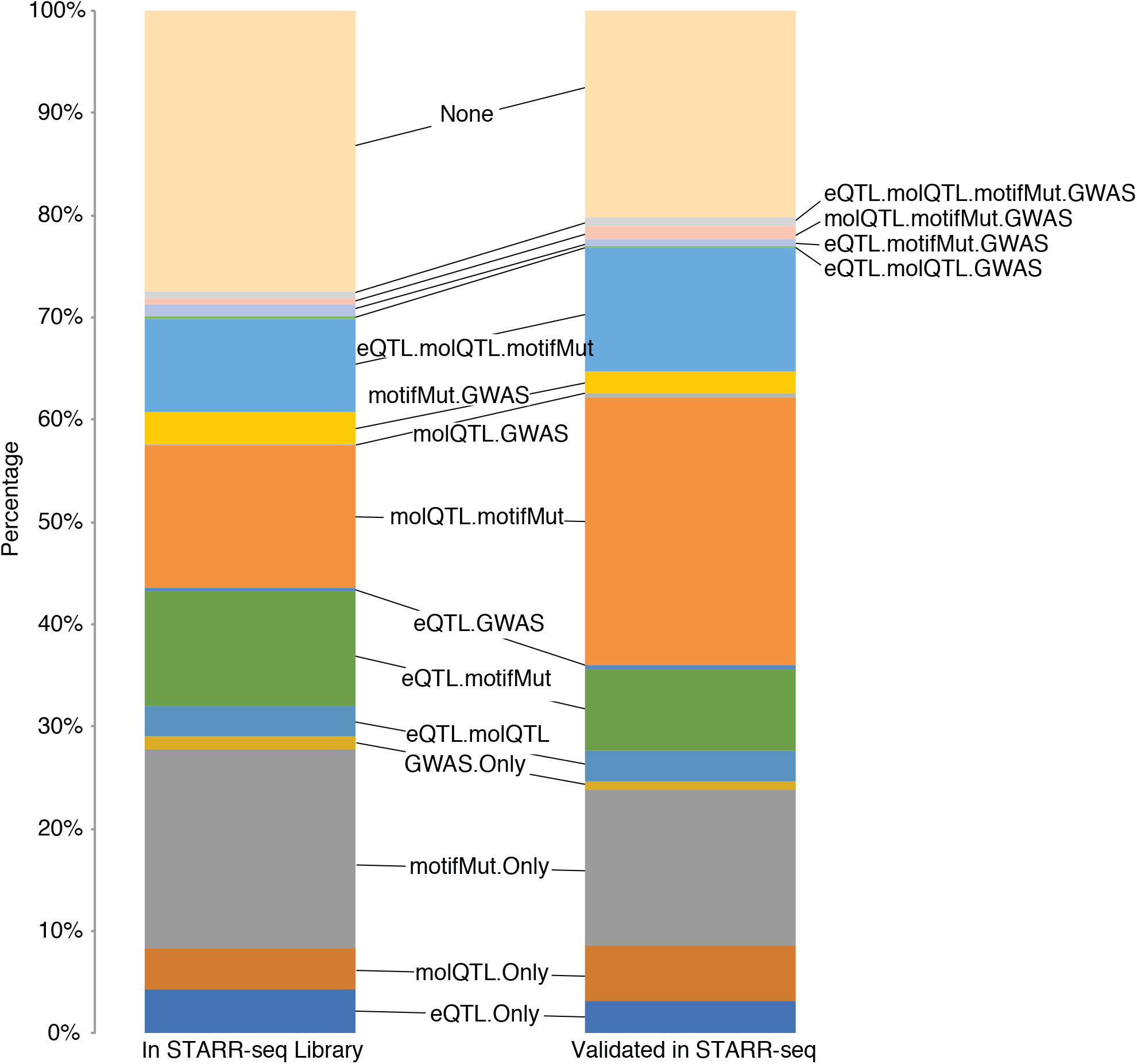
Contents of STARR-seq library and STARR-seq validated set. Stacked bar plot showing the proportion of the library and validated set of SNPs that had other layers of evidence (molQTL, eQTL, motif mutation, or GWAS)

**Figure S2.**
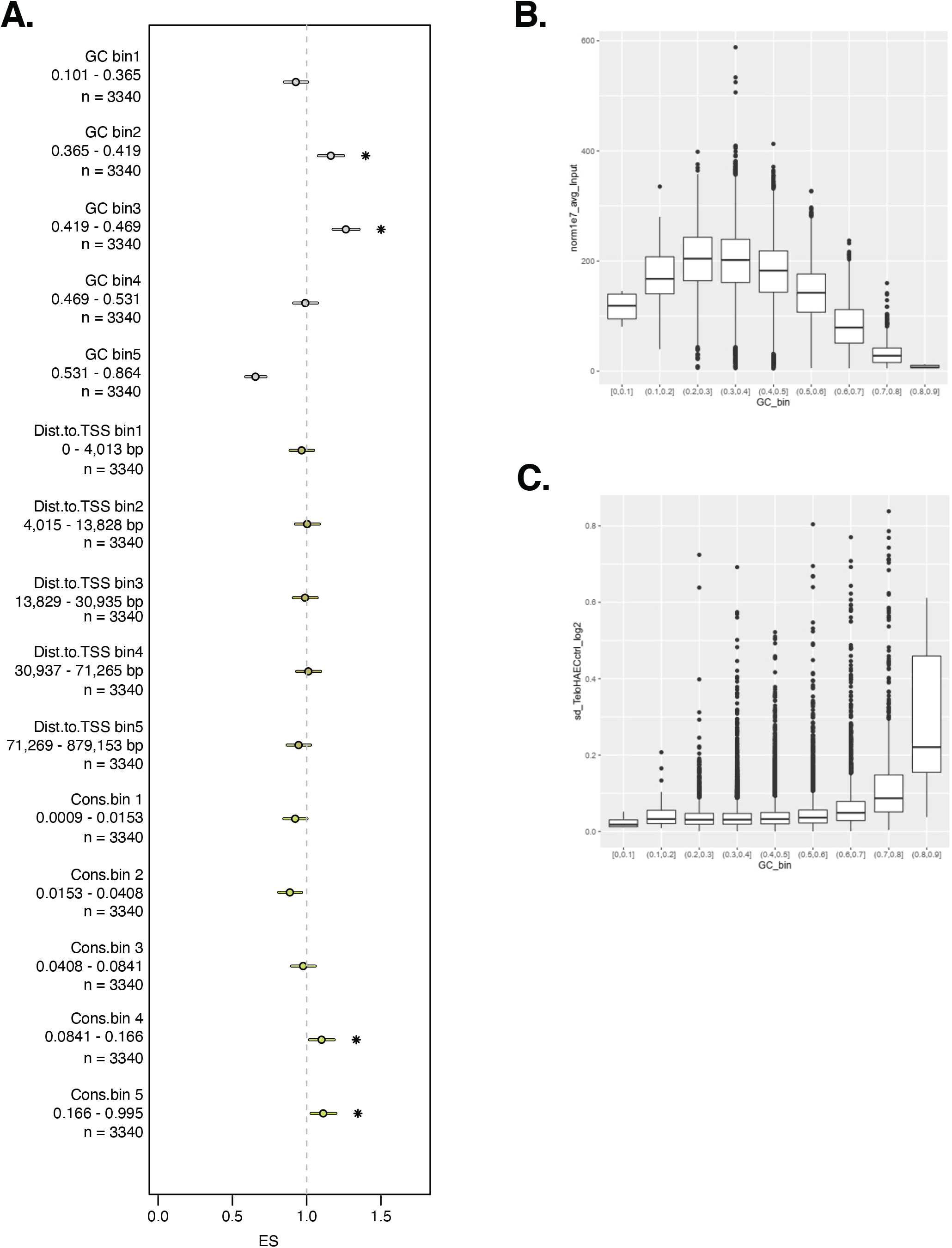
Genomic characteristic trends with validation and representation in the STARR-seq library. **A)** Enrichments of general genomic characteristics ranked by the characteristic and binned into equal number of SNPs. Characteristics included are average GC content of the DNA segments in the STARR-seq library that included the SNP, Distance to nearest TSS in the genomic context of the SNP, and average conservation of the regions surrounding the SNP that were input into the STARR-seq library. **B)** Oligo proportions in the plasmid DNA library as a function of GC%. **C)** Standard deviation of the replicates from library transfected TeloHAECs represented as RNA normalized to DNA.

**Figure S3.**
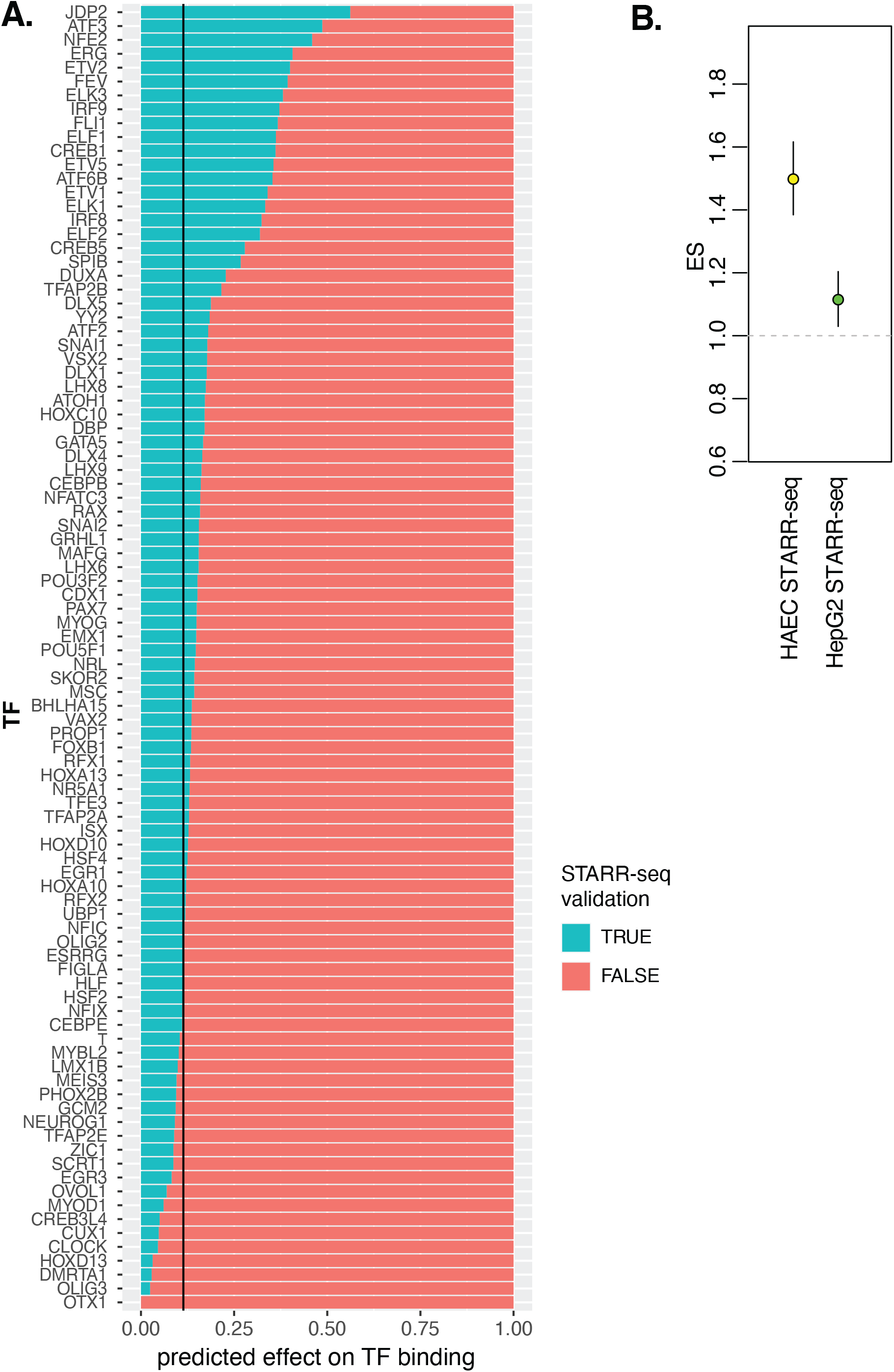
Allele specific enhancer function is associated with motif mutations and cell-type specific chromatin accessibility. **A)** The percentage of deltaSVM predictions (x-axis) that exhibited significant allelic effects in STARR-Seq. The 17% line signifies the average proportion of validated STARR-seq oligos accross the entire library. **B)** Enrichment of HAEC chromatin accessibility SNPs in STARR-seq performed in HAECs and STARR-seq performed in HepG2s.

**Figure S4.**
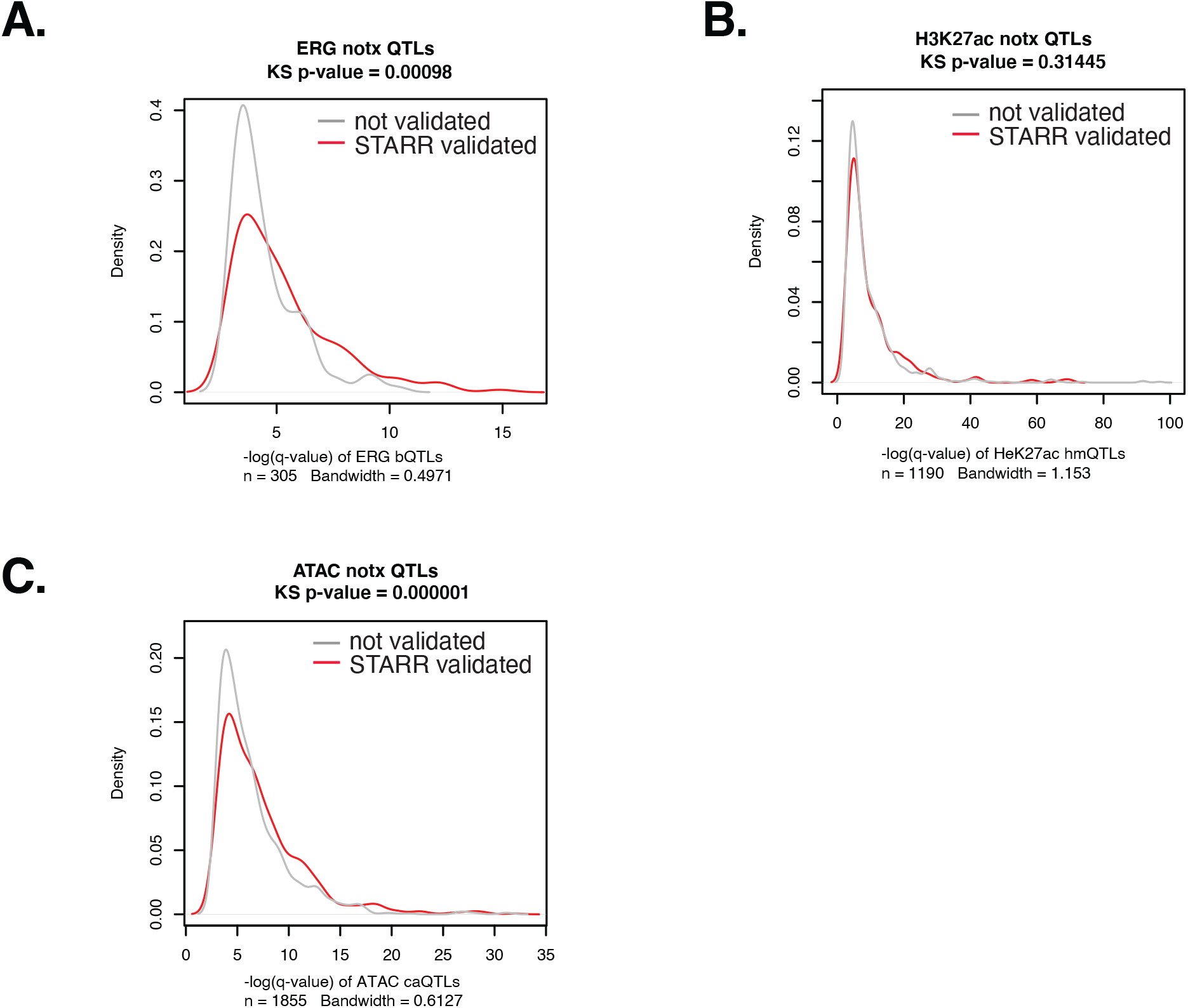
KS validation of enrichments observed for molQTLs in STARR-seq. Density plots of SNPs based on -log(q-value) in the molecular QTL analysis with grey being SNPs that did not validate in STARR and red being SNPs that did.

**Figure S5.**
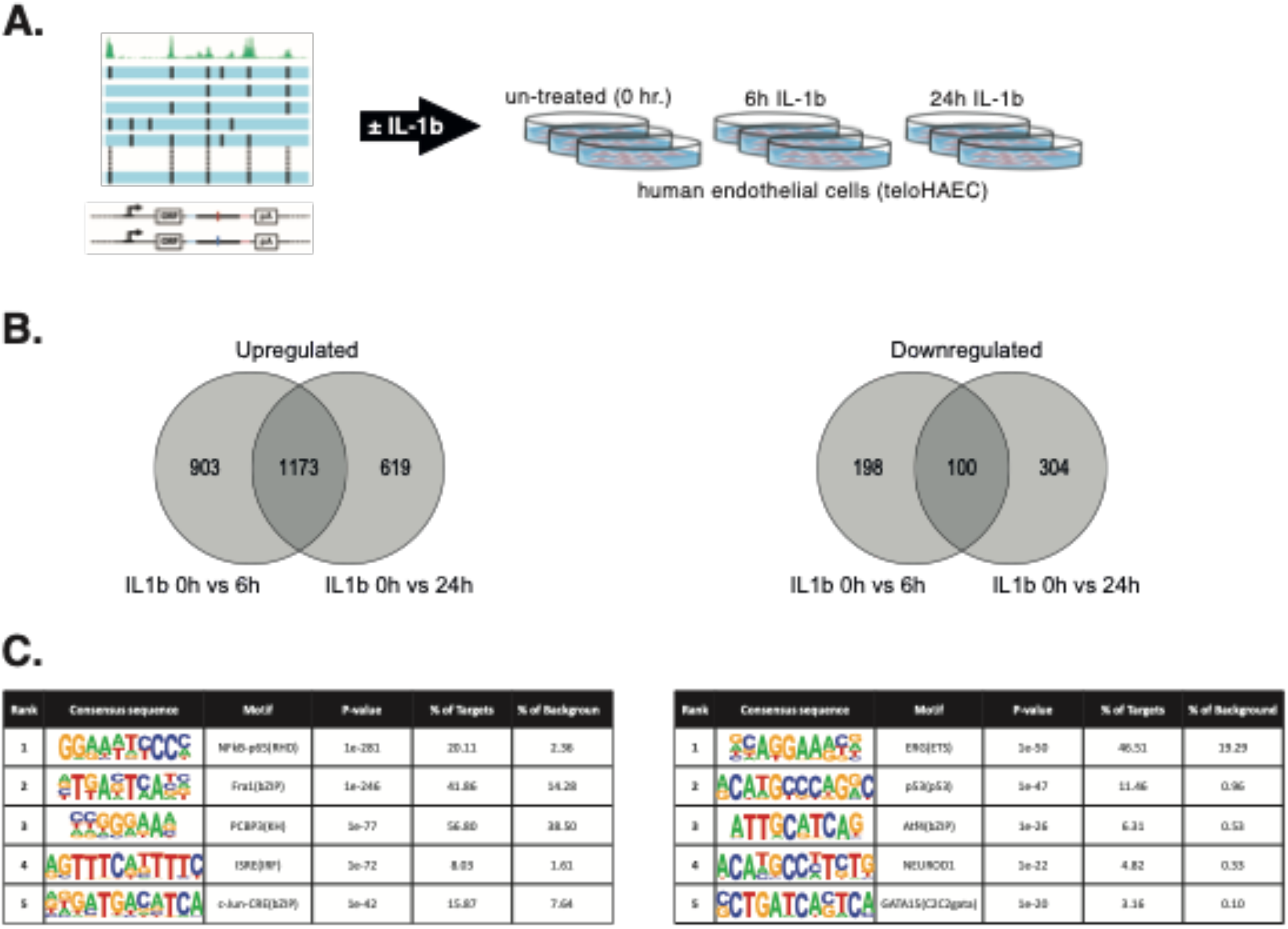
Analysis of the effect of inflammatory stimulus on STARR-seq enhancer activity in cultured human endothelial cells. **A)** Schematic of the experimental design **B)** Venn diagram of the oligos exhibiting differential enhancer activity in response to inflammatory treatment analyzed using DEseq2. **C)** *De novo* motif analysis of the oligos differentially expressed between control and interleukin-stimulated conditions.

**Figure S6.**
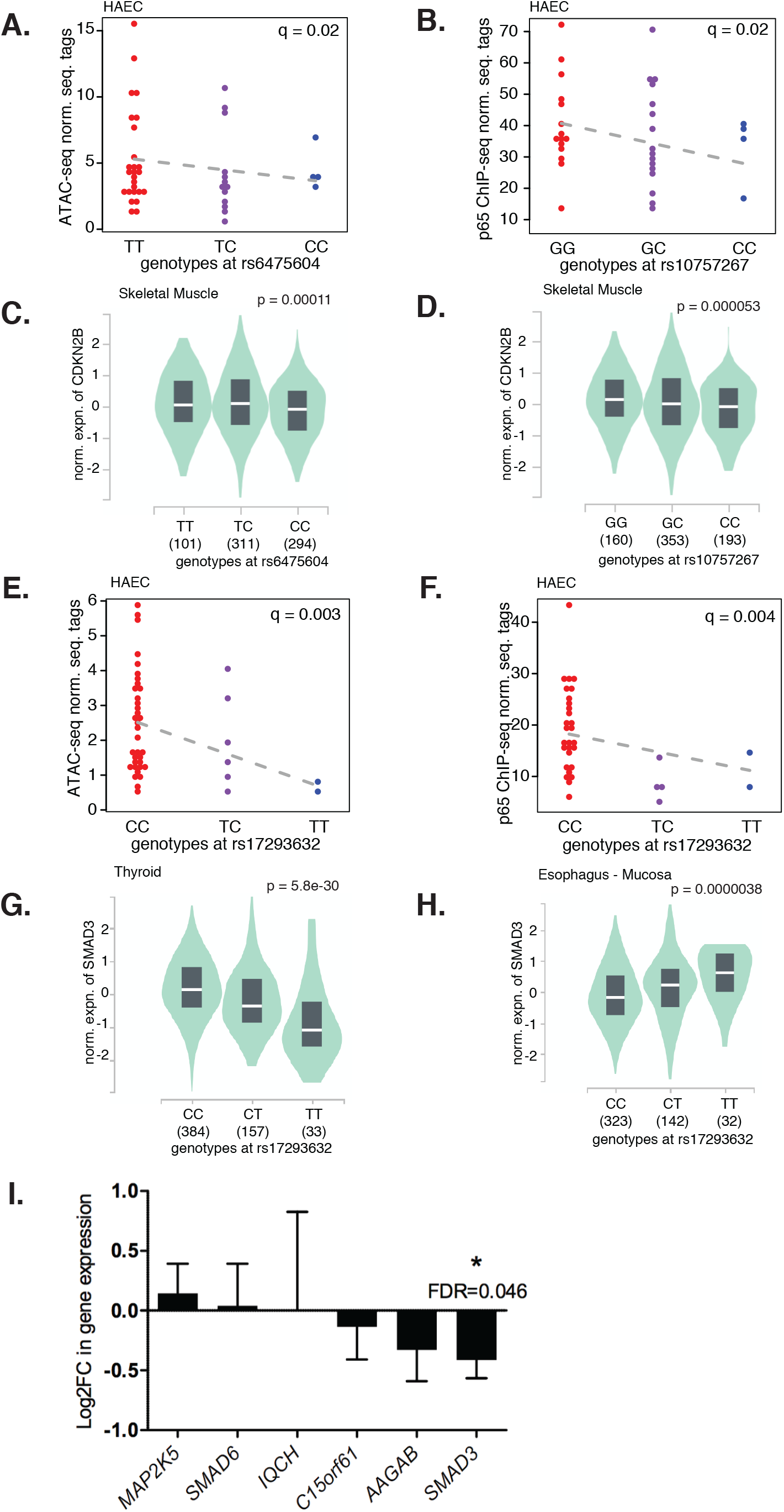
Coronary Artery Disease associated loci at genes CDKN2B and SMAD3. **A)** ATAC-seq tags in HAECs by genotypes of rs6475604 (RPKM normalized). **B)** p65 tags in HAECs compared between genotypes of rs10757267 (RPKM normalized). **C)** CDKN2B gene expression by genotypes at rs6475604 in Skeletal muscle from GTEx **D)** CDKN2B gene expression by genotypes at rs10757267 in Skeletal muscle from GTEx **E)** ATAC-seq tags in HAECs by genotypes at rs17293632 (RPKM normalized) **F)** p65 tags in HAECs by genotypes at rs17293632 (RPKM normalized) **G)** SMAD3 gene expression by genotypes at rs17293632 in Thyroid from GTEx **H)** SMAD3 gene expression by genotypes at rs17293632 in Esophagus - mucosa from GTEx **I)** Effect of rs17293632-carrying enhancer deletion on the expression of genes within 1 Mb. Only *SMAD3* demonstrates significant repression. DESeq2 was used to compare the enhancer deleted samples (n=4) to the controls (n=8). The data is represented as log2 fold change and standard error for the estimated coefficients on the log2 scale.

**Figure S7:**
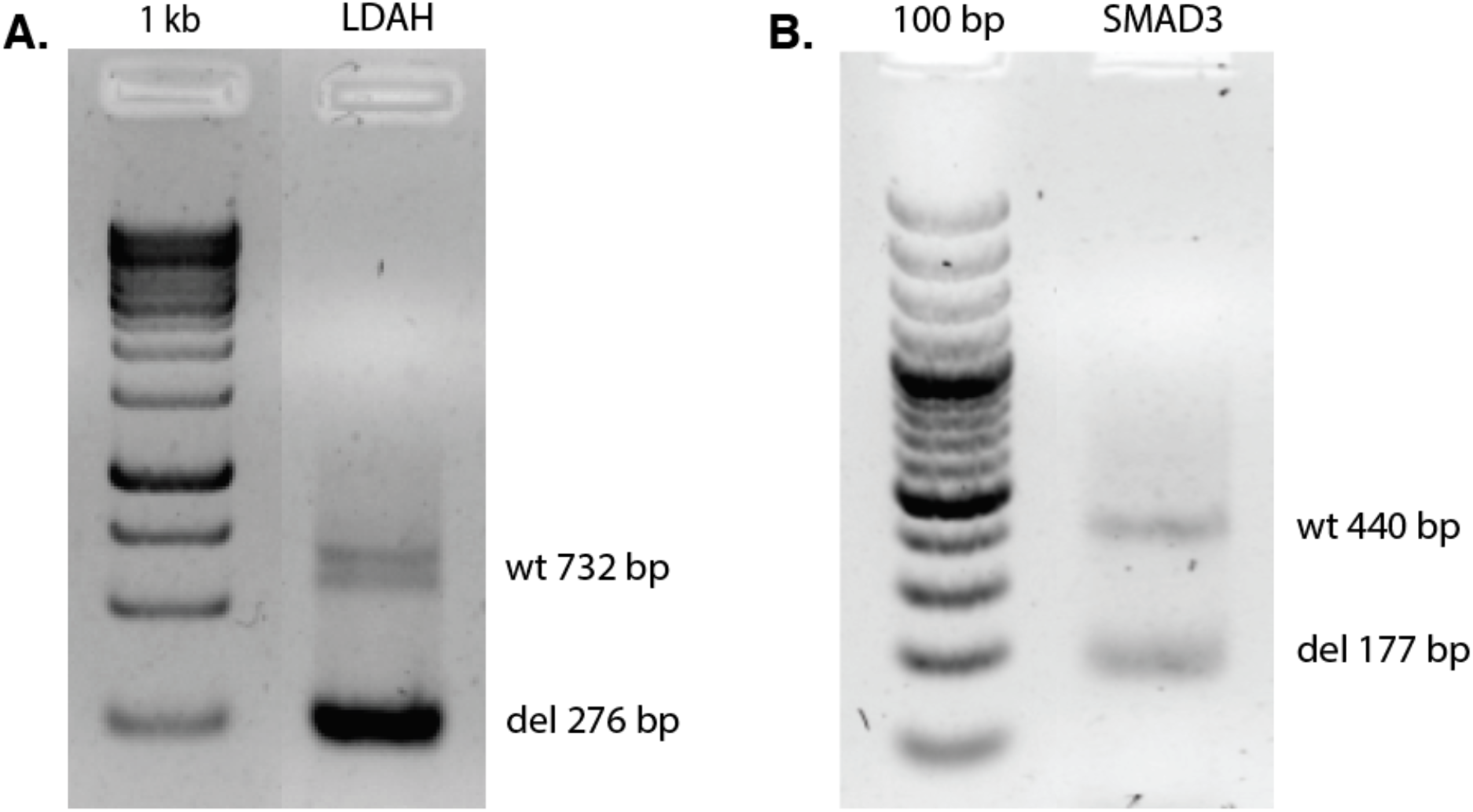
CRISPR-mediated perturbation of the validated endothelial enhancers. PCR analysis of the deletion efficiency of the enhancer in teloHAEC genomic DNA showing the deletion of enhancer regions predicted to regulate (**A**) LDAH and (**B**) SMAD3 genes.

**Table 1 –** Select list of credible functional non-coding SNPs

**Table S1 –** Expanded list of all credible functional non-coding SNPs

**Table S2 –** List of PCR primers and guide RNAs used for CRISPR/Cas9 deletion of target enhancers

